# The Where and When of Tracking Sound Sources Head-based auditory motion tracking integrates velocity and spatial error within a common delayed frame

**DOI:** 10.64898/2026.11.09.765311

**Authors:** Timo Oess, Siddhartha Krishna Iriventi, Marc O. Ernst

## Abstract

Auditory objects in everyday environments remain seldomly at the same position but move about in the environment and keeping track of their motion is an essential task for the auditory system to maintain a correct representation of auditory space/scene. Following such a moving sound source in two-dimensional space requires the nervous system to guide head movements using delayed and uncertain sensory information. How the auditory system integrates such information for combined horizontal and vertical movements is still debated.

We recorded head movements from 14 participants tracking a physically moving loudspeaker along unpredictable two-dimensional trajectories. Participants followed source motion with substantial delays and retained measurable spatial discrepancies. To explain these behaviours, we developed a hybrid model combining source velocity with the spatial discrepancy between source and head at the matched, delayed time points. The model predicts head velocity on previously unseen trajectories more accurately than formulations relying on either signal alone. Importantly, using preceding rather than current head orientation improved prediction, highlighting the importance of temporal correspondence between sensory and motor states. These findings suggest that head-based auditory motion tracking depends on complementary motion and spatial information represented within a common delayed reference frame.

## Introduction

The localization of stationary auditory objects has been investigated profoundly. However, typical auditory objects are moving about the enviroment and tracking their movement is less explored. Tracking a moving sound source requires the motor system to coordinate head movements using sensory information that arrives with delays of up to one second 3, 4], far longer than the *≈* 100 ms delays typical of visual smooth pursuit [6]. Despite this, listeners can follow a physically moving speaker with their heads across extended, unpredictable trajectories on the horizontal plane [4]. For such a special case, the auditory systems needs to integrate merely binaural cues like ITD and ILD. However how such a tracking is accomplished for complicated trajectories in two-dimensional space, requiring the additional integration of spectral cues remains unresolved.

Head-based auditory motion tracking poses two complementary requirements. First, head movements must follow the changing trajectory of the sound source: they must represent source *motion*, not merely a snapshot of source position. Second, head pointing must maintain spatial correspondence with the source: accumulated *positional discrepancy* must be corrected, even though both kinds of information—how the source is moving and where it is relative to the head— reach the listener only after the same processing delay. Auditory velocity coding is inherently imprecise: listeners underestimate the speed of moving sounds [9], velocity discrimination thresholds for acoustic targets are substantially higher than for visual ones [10], and auditory motion detection is better described by minimum spatial-displacement angles than by velocity thresholds [15]; together these suggest that a spatial correction term may carry particular weight in auditory tracking. In visual smooth pursuit, a hybrid integrating both signals outperforms either alone [5], but the analogous test has not been performed for head-based auditory motion tracking.

We continuously moved a physical loudspeaker along 100 randomised two-dimensional trajectories while 14 participants tracked it with their heads in darkness. Participants followed the global path of the source, but only with a delay of about one second and with a residual spatial error. Tracking therefore appears to rely on delayed information about where the source was and how it was moving (Figure 1A), which raises the question of which of these signals guides the head.

**Figure 1.**
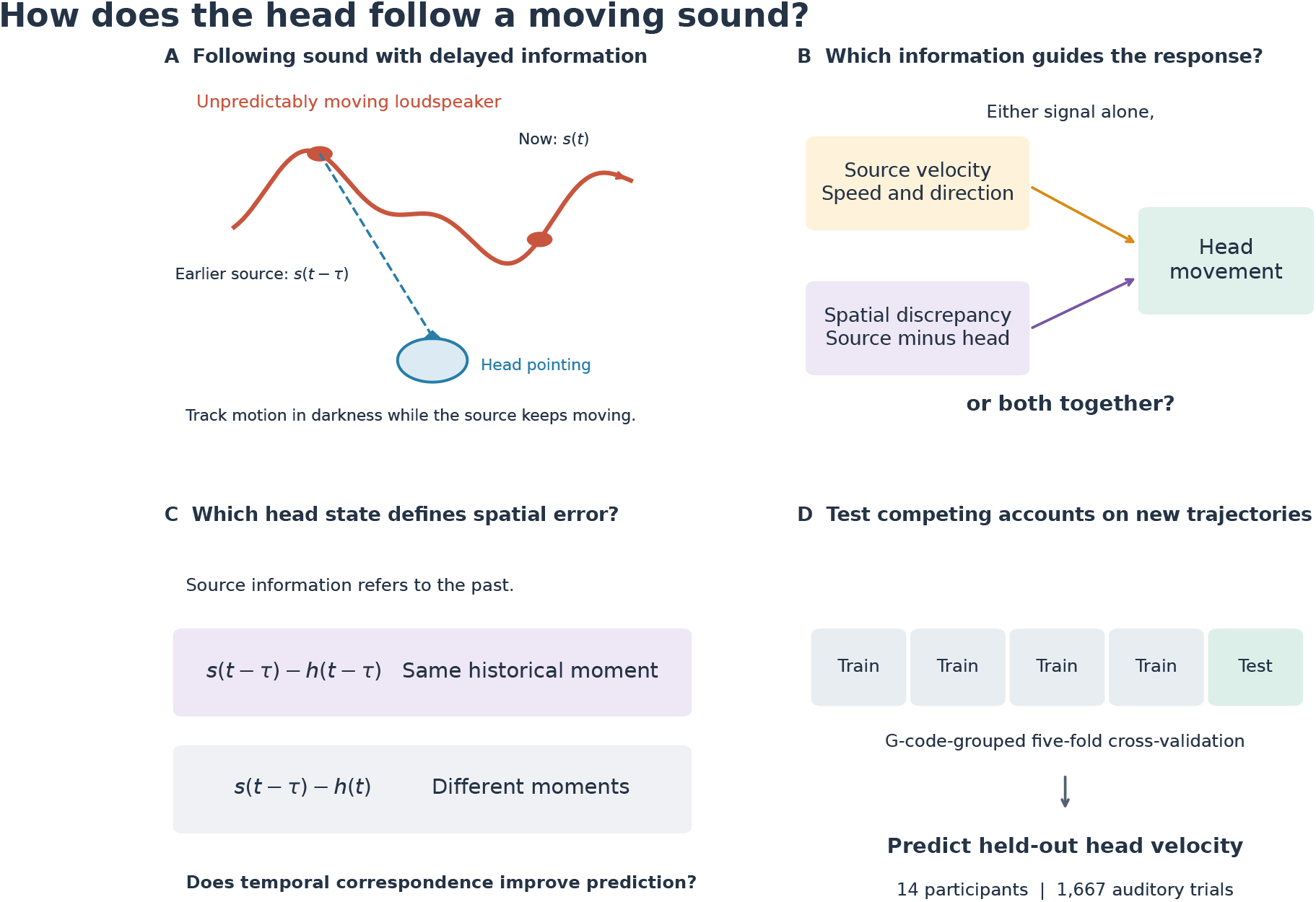
How does the head follow a moving sound using delayed information? **(A)** During head-based auditory tracking, the source continues to move while the listener responds to earlier sensory information; trajectory and head geometry are schematic. **(B)** Two candidate inputs could guide head velocity: source velocity and source–head spatial discrepancy. The study asks whether their combination predicts behaviour better than either input alone. **(C)** A second question concerns the time reference of the discrepancy: should the delayed source position be compared with the head position at the same earlier time, h(t − *τ*), or with the current head position, h(t)? **(D)** Competing models are evaluated on previously unseen G-code trajectories using grouped five-fold cross-validation (14 participants, 1,667 auditory trials). The diagram depicts the hypotheses and evaluation design, not an identified neural circuit.

Two candidate signals could guide such delayed tracking. A controller based on delayed source velocity can follow source motion but cannot correct positional drift. A controller based on the delayed source–head position difference can reduce spatial error but has no direct representation of source speed. A hybrid controller that combines both would meet both requirements outlined above (Figure 1B).

A second question concerns how the spatial error is computed. Because the auditory information is about one second old, it tells the listener where the source was at an earlier moment, while the head has kept moving since. The error can therefore be computed in two ways: by comparing that earlier source position with where the head points now, which mixes two different moments, or with where the head pointed at that same earlier moment, which keeps both in a common delayed frame (Figure 1C). Whether this distinction matters for predicting tracking behaviour has not been tested.

We hypothesised that head-based auditory motion tracking depends on both an estimate of source velocity and a representation of source–head spatial discrepancy, and that the temporally matched (historical) formulation provides the more informative spatial-error signal. To test whether these models *generalise* rather than merely overfit, all parameters were fitted on held-in source trajectories and evaluated on previously unseen ones (G-code-grouped five-fold cross-validation; Figure 1D; STAR Methods). We tested whether auditory-guided head movements are better predicted by a model that combines source motion with spatial discrepancy, and whether that discrepancy is most informative when source and head states are represented at the same historical time.

## Results

Participants sat in darkness in a sound-attenuated chamber, about 2 m in front of a 3 m *×* 3 m frontal plane across which a CNC robot moved a small loudspeaker. On each 20 s trial, the speaker followed one of 100 randomised two-dimensional trajectories at up to 0.25 m s^−1^ while playing white noise. Participants tracked the speaker by pointing their nose at it; head orientation was recorded by motion capture at 120 Hz (STAR Methods). The noise was either unmodulated or sinusoidally amplitude-modulated (AM) at 2, 10 or 18 Hz (80 % depth), giving four auditory conditions.

The amplitude modulation (AM) condition had no reliable effect on tracking accuracy or latency across the four auditory conditions (Friedman p = 0.28 for lag, p = 0.30 for vector correlation; exploratory residual-error difference did not survive independent cross-validation, Friedman p = 0.692; see Figure S2). All subsequent analyses pool across auditory conditions.

### Head-based auditory motion tracking shows consistent delay and spatial error

Head tracking showed a substantial positive sensorimotor lag. The median two-dimensional lag was 0.93 s (mean 1.00 s; Figure 2D), with a smaller lateral component (mean 0.77 s) and a larger vertical component (mean 1.37 s), consistent with the known asymmetry in auditory spatial resolution between azimuth and elevation [1, 2]. After applying each trial’s lag and removing a within-trial median pointing offset, the median residual spatial error was 0.238 m (Figure 2D), confirming that participants tracked the global trajectory of the source from a delayed and spatially imprecise estimate. Head velocity closely followed the temporal structure of source velocity after lag alignment, indicating that source motion—not merely source position—was an important determinant of tracking behaviour (Figure 2C; Figure S1).

**Figure 2.**
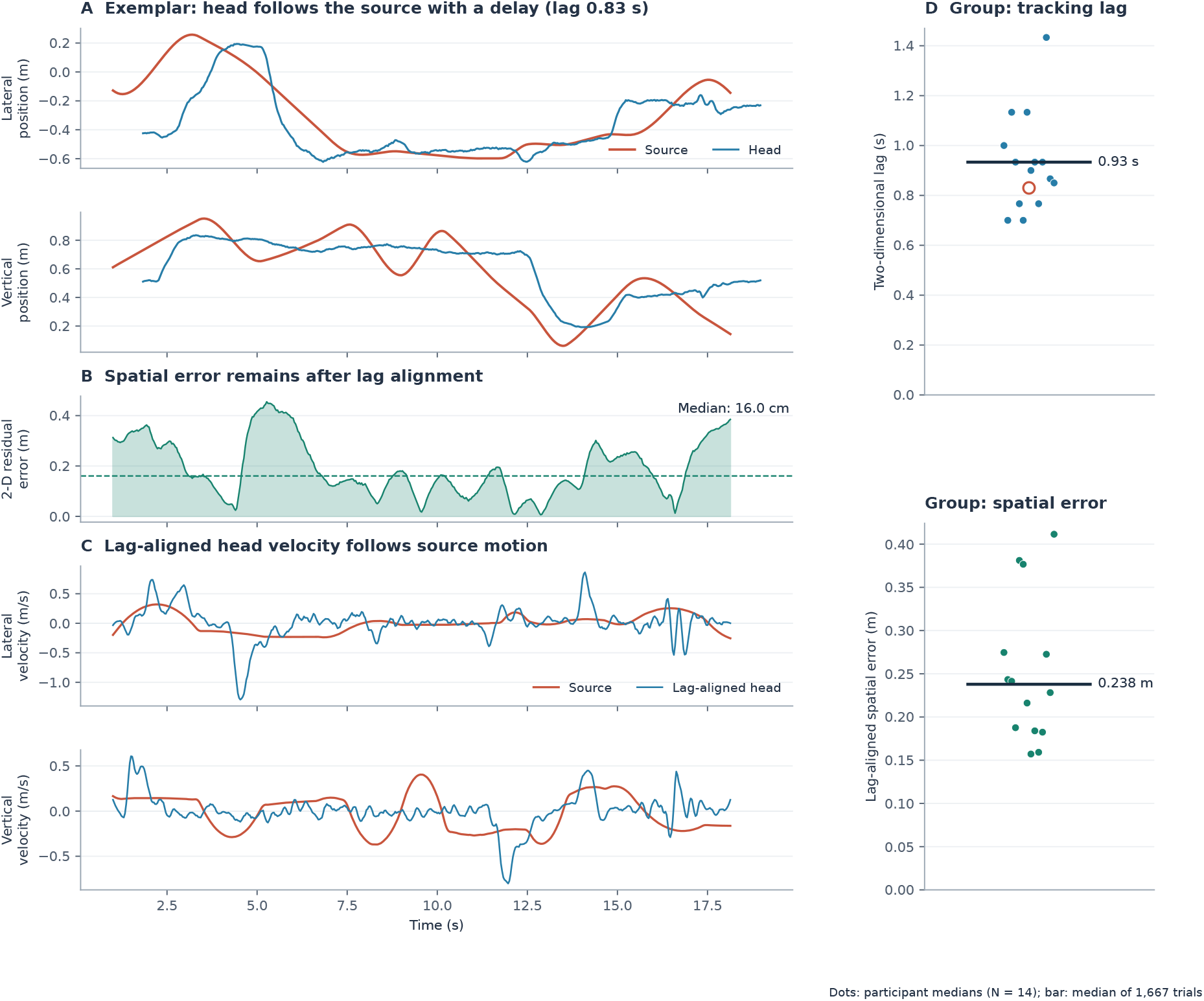
Behavioural characteristics of head-based auditory motion tracking. **(A–C)** Exemplar trial (participant D.H, white noise, G-code 70). **(A)** Lateral and vertical positions of the source and the head. Head traces are shown at their recorded time after removing the within-trial median offset; they lag the source by 0.83 s (two-dimensional cross-correlation lag). **(B)** Euclidean source–head residual after applying that lag and removing the offset; the median for this trial is 16.0 cm (dashed line). **(C)** Signed lateral and vertical velocities of the source and the lag-aligned head, differentiated on the 30 Hz grid with a third-order Savitzky–Golay filter (11 samples). **(D)** Group level: two-dimensional lag (top) and lag-aligned spatial error (bottom). Dots show each participant’s median across their auditory trials (N = 14); bars show the median across all 1,667 auditory trials (0.93 s and 0.238 m). The open circle marks the exemplar trial’s lag.

### A delayed hybrid controller captures source motion and spatial discrepancy

Head-based auditory motion tracking exhibited two consistent features (Figure 2): movements followed the temporal evolution of source motion despite substantial delays, yet retained a measurable spatial discrepancy from the source. These observations motivate a model with two complementary inputs. In brief, the model predicts how fast and in which direction the head moves at each moment from two signals that both refer to the same earlier moment, one delay *τ* ago: how the source was moving then, and how far the head was from the source then. Each signal is scaled by its own gain, and the two contributions are added.

Two input signals operate within this delayed framework. Head movements closely track source motion, motivating a source-velocity term scaled by a gain matrix G, 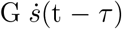 representing the source’s speed and direction at the earlier time t − *τ*. Spatial discrepancy between source and head persists after lag alignment (each trial shifted by its own cross-correlation lag and corrected for a constant pointing offset; STAR Methods), motivating a corrective term proportional to the position difference at the same delayed time, K [s(t − *τ*) − h(t *− τ*)]. Combining these two terms with a participant-level velocity intercept b (a constant drift of the head that does not depend on the source) gives the delayed hybrid model (Figure 3A):

**Figure 3.**
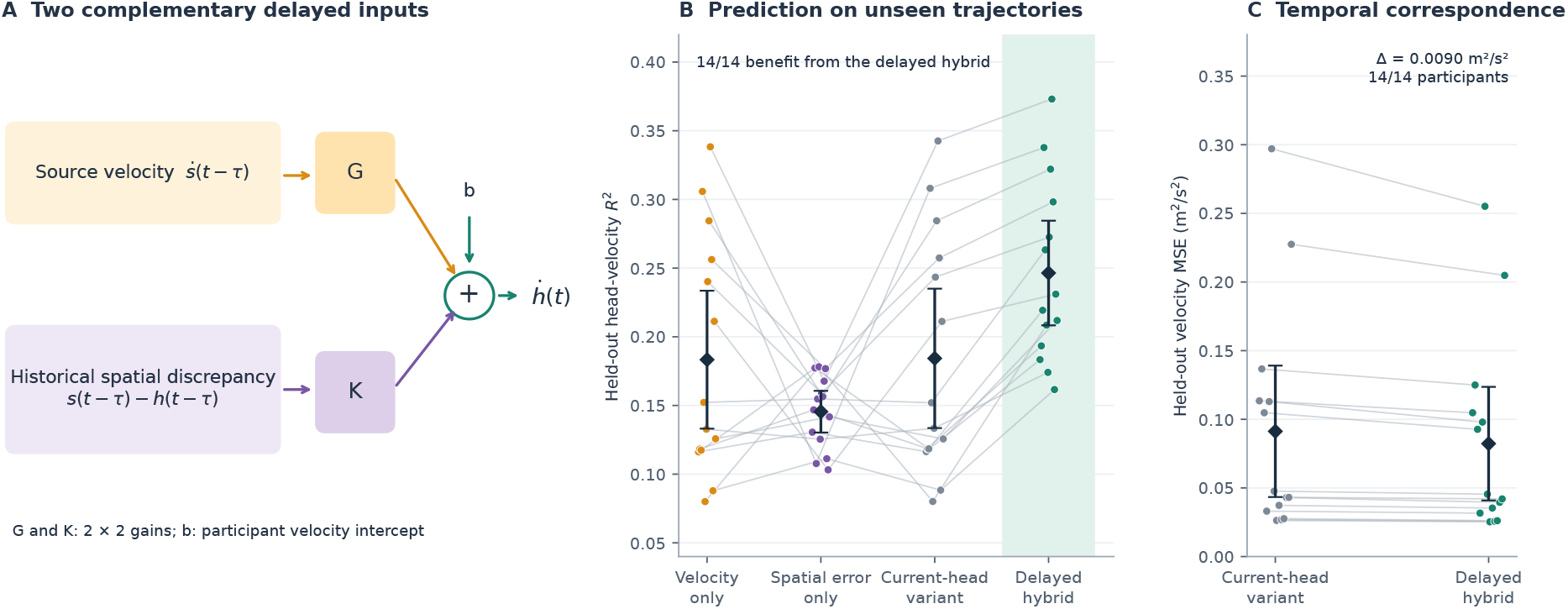
A delayed hybrid model combines motion and spatial information, and temporal correspondence improves prediction. **(A)** Source velocity and source–head spatial discrepancy enter parallel pathways, each scaled by its own gain matrix (G, K). Both source and head positions in the discrepancy are evaluated at t − *τ*. Their contributions and a participant-level intercept b predict head velocity (Equation 1); this is a statistical model diagram. **(B)** Held-out head-velocity R^2^ for the four competing models, averaged across five G-code-grouped folds per participant and evaluated on common test windows. The delayed hybrid model improves prediction over both single-input variants and the current-head variant in all 14 participants (one-sided Wilcoxon p = 6.1 × 10^−5^ for each comparison). **(C)** Held-out head-velocity MSE for the current-head variant, which pairs the earlier source position with the current head position, and the delayed hybrid model, which compares both at the same earlier time. The delayed hybrid model reduces mean MSE by 0.009 m^2^/s^2^ (one-sided Wilcoxon W = 105, p = 6.1 × 10^−5^; 14/14 participants). In B and C, grey lines connect each participant’s values (N = 14); dark diamonds and bars show the group mean and t-based 95 % confidence interval.

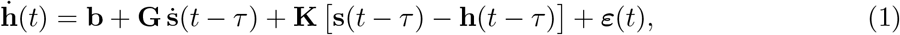

where h(t) and s(t) are two-dimensional head-pointing and source positions, b is the participant-level velocity intercept, and G, K are 2*×*2 gain matrices. Setting K = 0 yields the velocity-only variant; setting G = 0 yields the spatial-error-only variant. A current-head variant replaces h(t *− τ*) with h(t), testing whether temporal correspondence within the discrepancy term matters.

The median cross-correlation lag of *≈* 0.93 s (Figure 2; Figure S1) reflects the combined effects of sensory processing, motor planning, and movement dynamics; we use it to constrain the range over which the model delay *τ* is selected, with the optimal *τ* estimated independently from the fitting data.

### Both signals are required for accurate prediction on unseen trajectories

To fit our model and subsequently test it, we split the trajectories (g-codes) into 5 different groups each containing 20 trajectories. Models were fitted on four G-code groups and tested on the held-out fifth group (five-fold, G-code-grouped cross-validation; STAR Methods). Because each model selects its own delay *τ*, a model with a longer delay cannot predict the first *τ* seconds of a held-out trial. To compare all models on exactly the same data, every model in a fold was therefore scored only on samples after the longest delay selected in that fold.

The delayed hybrid model (Equation 1) explained R^2^ = 0.25 of the variance in held-out head velocity (mean MSE = 0.0822 m^2^/s^2^; we evaluate velocity because the model specifies how the head moves, whereas head position is dominated by slow components that every model captures), predicting head velocity on previously unseen source trajectories more accurately than either component alone. Removing the velocity term (spatial-error-only variant, R^2^ = 0.15, MSE = 0.0900 m^2^/s^2^) reduced explained variance by 0.10 (W = 105, p = 6.1*×*10^−5^, 14/14 participants). Removing the discrepancy term (velocity-only variant, R^2^ = 0.18, MSE = 0.0913 m^2^/s^2^) reduced it by 0.06 (W = 105, p = 6.1*×*10^−5^, 14/14 participants). The two restricted variants did not differ significantly from each other (W = 49, p = 0.596), demonstrating that source velocity and spatial discrepancy provide complementary information for predicting auditory-guided head movements (Figure 3B; Figure S5).

### Temporal correspondence improves prediction of auditory motion tracking

During delayed auditory pursuit, the source continues moving while the listener’s head responds to earlier sensory information: with a lag of about one second (Figure 2A), the source has moved on by the time the head reacts to it. Consequently, the head’s present orientation and its orientation when the relevant source information was available can differ substantially. A controller based on current head position and one based on preceding head position therefore make different predictions, even when both use the same delayed source trajectory. Comparing these formulations allows us to examine which temporal representation better accounts for observed tracking behaviour.

A current-head variant that replaced h(t *− τ*) with the current head state h(t) achieved R^2^ = 0.18 (MSE = 0.0912 m^2^/s^2^), not reliably different from the pure variants. The delayed hybrid model explained a greater proportion of variance (ΔR^2^ = 0.06, ΔMSE = 0.009 m^2^/s^2^; W = 105, p = 6.1*×*10^−5^, 14/14 participants). The delayed-state formulation is therefore the better-performing predictive representation: temporally matched source and head information is more informative for predicting tracking behaviour than pairing delayed source estimates with the current head state (Figure 3C).

### The contributions of the two signals vary with trajectory dynamics

The delayed hybrid model uses the same gains G and K throughout. Even so, the share of each term in the predicted head velocity is not fixed, because it also depends on the size of its input: when the source moves fast, the velocity input is large and the velocity term contributes more; where the path bends, the source–head discrepancy tends to grow and the spatial-error term contributes more. We quantified this as the velocity-term share, the magnitude of the velocity term divided by the summed magnitudes of both terms (STAR Methods, Trajectory-regime attribution). In held-out samples, the velocity-term share increased with source speed (from 0.41 in the slowest quintile to 0.59 in the fastest) and decreased with source curvature (from 0.54 to 0.43; Figure S6). Because the gains are constant, these shifts reflect how the inputs vary along the trajectory, not an adaptive re-weighting of the two signals. Unlike the primary model comparison, this attribution analysis used trial-based rather than G-code-grouped folds, so it is descriptive. A model variant whose gains change with source jerk is reported in Supplementary Note S1.

## Discussion

Listeners tracked unpredictable two-dimensional source trajectories with their heads, but with a long delay and a residual spatial error. The median lag of 0.93 s is far longer than the *≈* 100 ms delays of visual smooth pursuit [6] and longer than the head-pursuit lags of a few hundred milliseconds reported for horizontal sound motion [4]. Vertical tracking lagged lateral tracking by about 0.6 s (1.37 vs 0.77 s), in line with the poorer resolution of elevation, which depends on spectral rather than binaural cues [1, 2]. After the lag was removed, the head still missed the source by about 24 cm, roughly 7° at the 2 m listening distance. Amplitude modulation of the noise changed neither lag nor accuracy, so added envelope fluctuations at the rates tested did not aid tracking. Together, these results indicate that listeners follow a moving sound from a delayed and spatially imprecise estimate of its position and motion, which is what motivated the model.

Head-based auditory motion tracking is more accurately predicted by a delayed hybrid model that combines source velocity with historical source–head spatial discrepancy than by either signal alone. This result held across all 14 participants on previously unseen trajectories, establishing that both inputs provide complementary predictive contributions. Two properties define the better-predicting model: it is *hybrid* (combining velocity and spatial-discrepancy signals) and it evaluates spatial error in a *temporally matched frame* (comparing source and head positions from the same earlier time).

The temporal correspondence result is the clearest mechanistic implication of the model. When auditory information about source position arrives substantially delayed, a tracking system that computes the source–head discrepancy from a temporally matched reference—both quantities evaluated at t *− τ* —has a more coherent spatial reference than one that must bridge the delayed source estimate with the head’s current position. The spatial-error term s(t *− τ*) − h(t *− τ*) represents the discrepancy as it stood when the relevant sensory information was available; the current-head formulation s(t *− τ*) − h(t) conflates source position at one moment with head position at another [8]. Although this result is consistent with a shared delayed internal representation of source and head position, it does not establish that the nervous system relies exclusively on historical state information; current proprioceptive and efference-copy signals may contribute to both representations. The account makes a testable prediction: if the error is computed in a delayed frame, an unexpected perturbation of the head should change the corrective response only after the delay *τ*, whereas a controller that uses the current head orientation would respond to it at once.

Visual smooth pursuit integrates retinal velocity and retinal position error, with their relative weight adapting to target dynamics [5, 6, 11, 12, 17]. The present results mirror this hybrid architecture: both velocity and spatial-discrepancy information are required for accurate prediction on unseen trajectories, and neither signal alone accounts for tracking behaviour. These findings reveal a computational similarity to visual smooth pursuit, in which motion and spatial-error information also contribute to tracking. However, head movements operate over substantially longer behavioural delays—on the order of one second—making the temporal correspondence between source and head representations particularly relevant.

These findings extend those of García-Uceda Calvo et al. [4], who showed accurate head pursuit of sounds moving unpredictably in the horizontal plane. Their responses were well described by a second-order low-pass filter with a short pure delay (10–98 ms), with head lags of a few hundred milliseconds. Position-error and velocity-error regressors described their data about equally well, so velocity tracking and rapid position sampling could not be told apart. Three differences stand out. First, our lag was roughly three times longer than in their example listener.

Our trajectories were two-dimensional, so elevation had to be estimated from spectral cues. Our trajectories were also slow (*≤* 0.25 m s^−1^, about 7° s^−1^ at 2 m), which permits long lags without large errors; the measures also differ, because our cross-correlation lag includes movement dynamics that their model assigns to the filter rather than to the pure delay. Second, by testing predictions on held-out trajectories we could separate the two signals: both contributed, and neither sufficed alone. Third, they found that damping increased across trials, an adaptation to the shared spectrum of their stimuli; our randomised trajectories had no common spectrum and our gains were fixed, so the trajectory-dependent shifts in Figure S6 reflect input covariation, not adaptation.

The delayed hybrid model identifies predictive relationships between source dynamics and head movements; it does not uniquely determine the neural implementation. The fitted delay and gain structure is consistent with a forward-model account in which the motor system predicts the sensory consequences of its own commands to compensate for feedback latency [13, 14, 18], but other neural architectures could produce equivalent input–output relationships. The velocity-term share analysis (Results; Figure S6) shows that even with constant gains, the relative weight of the velocity and spatial-error terms varies systematically with trajectory statistics: the velocity term dominates at high source speed [16] and the spatial-error term at high curvature. These are input-covariation patterns, not validated gain-scheduling effects. Disambiguating the two requires designs that independently perturb source velocity and source–head position error at known trajectory phases. A jerk-dependent-gain extension is described in the Supplementary Information (Supplementary Note S1).

In conclusion, listeners track a sound moving in two dimensions with a delay of about one second and a residual spatial error. Their head movements are best predicted when source velocity and source–head spatial error are combined, and when that error compares source and head at the same earlier moment. Head-based auditory motion tracking thus appears to rely on motion and position information held together in a common delayed reference frame.

## STAR Methods

### Resource Availability

#### Lead contact

Further information and requests for resources should be directed to Timo Oess.

### Data and code availability

Raw motion recordings (1,737 CSV files), analysis scripts (Python 3.10), and result tables will be deposited on Zenodo upon acceptance.

### Experimental Model and Study Participant Details

Fifteen participants (ages 23–28 years) took part. One participant (A.P.) did not complete the full set of conditions required for a balanced within-subjects design and was excluded, yielding N = 14 (1,737 recordings; 1,667 randomised auditory trials after excluding 70 visual familiarisation trials). Written informed consent was obtained; the study followed the Declaration of Helsinki and was approved by the Ulm University ethics committee (EG-IIP-2025089).

### Method Details

#### Apparatus and stimuli

The experiment was conducted in a sound-attenuated chamber. A small loudspeaker was mounted on a CNC robot that moved it across a 3 m *×* 3 m frontal plane *≈* 2 m in front of the participant. Drive motors were housed in a separate room. The speaker followed 100 randomised (x, z) trajectories encoded as G-code programmes at a capped speed of 0.25 m s^−1^.

Head orientation was recorded at 120 Hz by OptiTrack motion capture via a rigid marker array on head-tracking goggles. Reflective markers were also mounted on the speaker carriage.

Each auditory trial used 20 s white noise (65 dB SPL), either unmodulated or sinusoidally amplitude-modulated at 2 Hz, 10 Hz, or 18 Hz at 80 % depth (carrier amplitude reducing to 20 % of unmodulated level at the trough; m = 0.8). RMS level was equated across conditions. Visual familiarisation trials (5 per session) used a visible speaker and are excluded from all quantitative analyses.

#### Signal preprocessing

Raw CSV files contained synchronised speaker (id = 1) and head-pointing (id = 2) position streams. For each trial, streams were merged on timestamp, sorted, and converted to seconds. The first and last second of each recording were excluded to remove cueing and endpoint behaviour. Signals were linearly interpolated to a common 30 Hz grid. Position was differentiated using a third-order Savitzky–Golay filter [7] with a 0.35 s window; velocity, acceleration, and jerk were the first, second, and third derivatives. Curvature was computed as

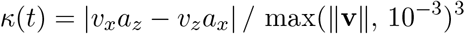

#### Lag-aligned spatial precision

For each trial, the source and head signals were cross-correlated over signed lags from −1 to +4 s using overlapping samples only (no zero padding). The selected lag maximised the mean of the and z-axis correlations. After applying this lag, a constant two-dimensional median offset was removed within the trial. The Euclidean norm of the remaining residual gave lag-aligned spatial error.

For the AM analysis, the grand-mean residual for each G-code was subtracted from every corresponding trial (G-code-adjusted residuals). Participant-level means were compared across four auditory AM conditions using Friedman tests; pairwise comparisons used Wilcoxon signed-rank tests with Holm correction.

#### Control-law model comparison

The Savitzky–Golay head-velocity estimate and position signals were decimated to 10 Hz. Source velocity was computed by numerical differentiation of decimated source coordinates. The primary model was evaluated against three restricted variants and one exploratory extension. The primary model is the delayed hybrid (Equation 1). Three restricted variants served as ablation controls:

- **Velocity-only variant**: 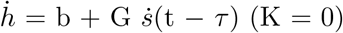
- **Spatial-error-only variant**: 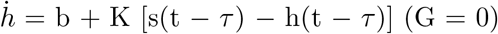
- **Current-head variant**: 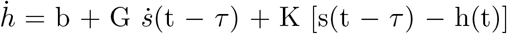

A jerk-dependent extension is described in Supplementary Note S1.

Candidate delays ranged from 0 to 3 s in 0.1 s increments. For every candidate delay, model parameters (b, G, K) were estimated by ordinary least squares over both output dimensions jointly. The delay minimising the MSE on the fitting data was selected independently for each model and fold.

#### G-code-grouped five-fold cross-validation

G-codes were assigned deterministically to one of five folds so that no source trajectory in any test fold appeared in fitting. Parameters and delays were fitted on four G-code groups. Within every test fold, all primary models were then evaluated on the *identical* held-out time samples beginning at the largest delay selected during fitting in that fold (the common-window constraint), ensuring strictly fair comparison. The primary outcome was held-out head-velocity MSE in m^2^/s^2^. To aid interpretation, R^2^ = 1 MSE / Var(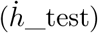) was also computed, where Var(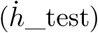) is the mean per-dimension variance of the held-out head velocity window; reported R^2^ values are participant means across folds. Participant means across held-out folds were the unit of statistical inference; comparisons used Wilcoxon signed-rank tests. Four trajectory-feature gain tests were Holm-corrected

#### Trajectory-regime attribution

For each held-out sample, the squared two-dimensional prediction error of the velocity-only and spatial-error-only models was compared to assign a pure-model win. Hybrid contribution shares were computed from the Euclidean magnitudes of 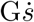 and K(s − h), excluding the intercept. Within each participant, samples were divided into quintiles of speed, acceleration, jerk, and curvature. Quintile profiles are reported descriptively, without inferential tests. Unlike the primary model comparison, this analysis assigned trials to folds by trial index (trial i to fold i mod 5) rather than by G-code, so trajectories could recur across fitting and test data; it is therefore descriptive.

#### High-jerk event analysis

High-jerk events were sustained top-decile source-jerk periods lasting at least 0.2 s; one local maximum was retained per event with a minimum separation of 0.5 s. Event-related residual error was measured in a 0.2–0.8 s post-event window and compared with adjacent pre/post baseline windows. Low-jerk pseudo-events (sustained bottom-quartile source-jerk periods) from the same trials served as within-trajectory controls.

#### Quantification and Statistical Analysis

All statistical tests used N = 14 participants as the unit of inference; no individual time sample was treated as independent. Wilcoxon signed-rank tests were used throughout (non-normal distributions expected due to the few participants near the performance ceiling). Holm correction was applied to families of four trajectory-feature tests. Analyses used Python 3.10 with NumPy 1.26, SciPy 1.11, pandas 2.1, scikit-learn 1.3, and tqdm.

