## Supplementary for "The Where and When of Tracking Sound Sources Head-based auditory motion tracking integrates velocity and spatial error within a common delayed frame"

### Figure S1. Cross-Correlation Curves

Cross-correlations between speaker and head-position time-series confirmed consistent auditory lags across conditions (mean lateral: 0.77 s; mean vertical: 1.37 s; mean 2D: 1.00 s) and a substantially shorter visual lag (mean ≈ 0.08 s; Figure S1). No condition effects on peak correlation or lag were detected across the four auditory AM conditions.

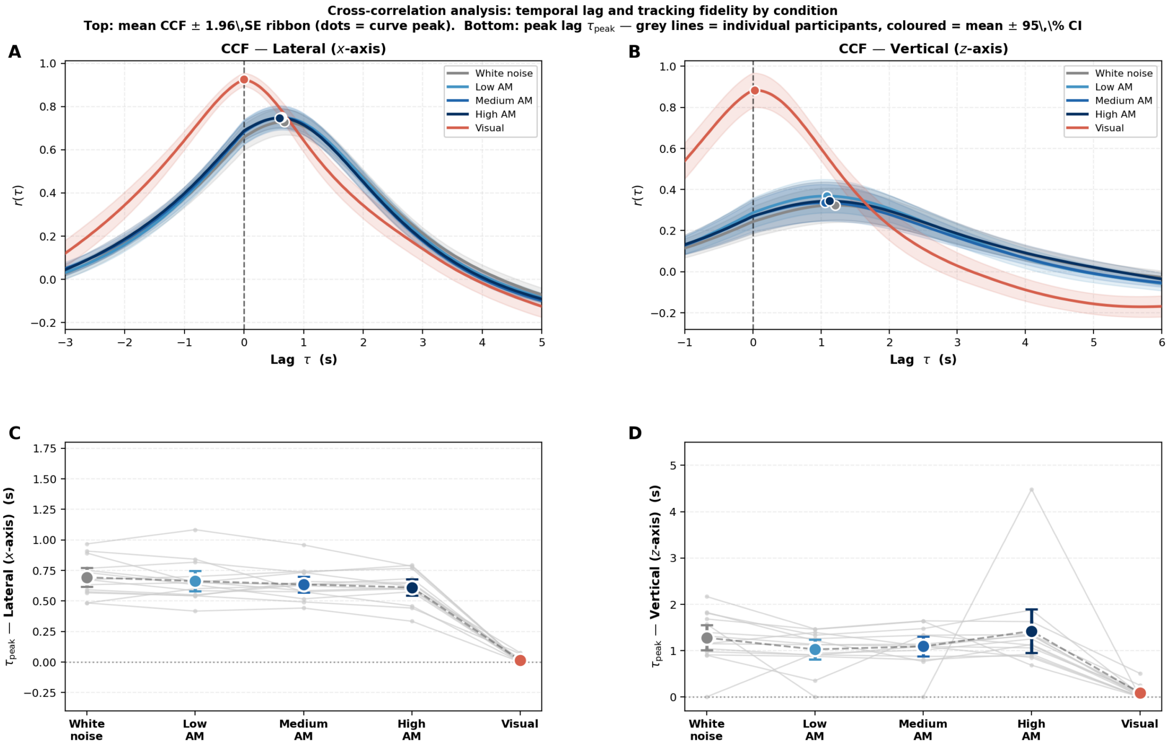


Figure S1: **Cross-correlation analysis of source and head position.** Mean Pearson-normalised cross-correlation functions for lateral and vertical coordinates, with ±1.96 SE ribbons and peak markers, together with participant-level peak lags by stimulus condition. The visual familiarisation condition is shown as a descriptive benchmark and is excluded from the auditory comparisons.

### Figure S2. Exploratory Low-AM Accuracy Effect

An exploratory trial-aligned residual analysis suggested lower residual error for the 2 Hz AM condition. The effect did not survive independent cross-validation: when lag and offset were estimated from the fitting G-codes only and applied unchanged to held-out trials, the low-AM versus white-noise G-code-adjusted difference was 0.013 m (Friedman p = 0.692; Wilcoxon p = 0.391).

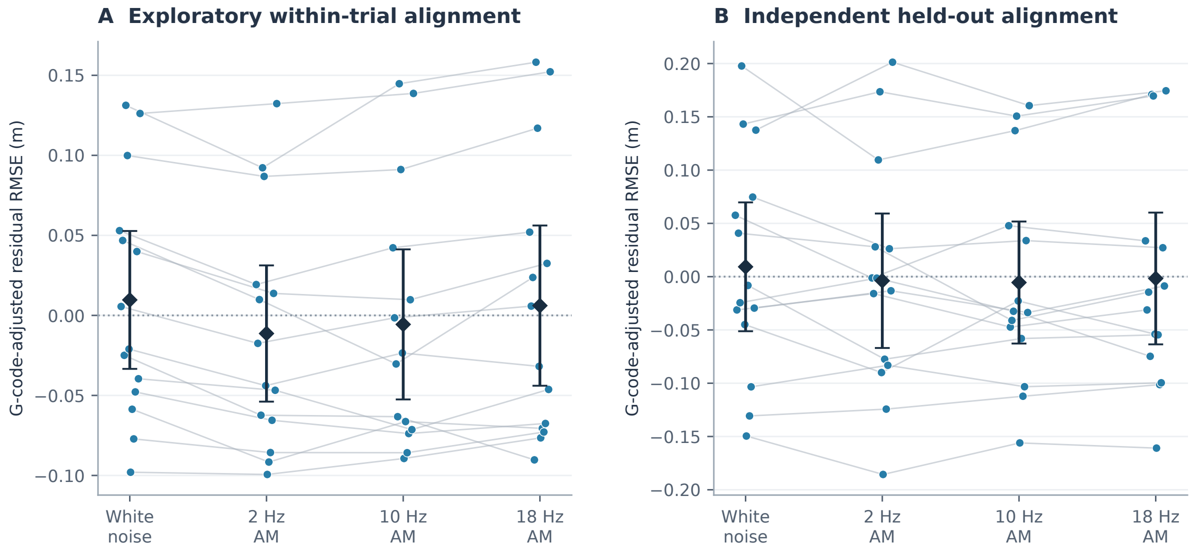


Figure S2: **Amplitude-modulation comparisons depend on independent alignment. (A)** Exploratory residual RMSE after trial-specific lag and offset estimation, adjusted by subtracting the across-participant mean for each G-code and averaged within participant and condition. **(B)** Held-out residuals obtained when each participant's lag and offset are estimated from the fitting trajectories and applied unchanged to unseen trajectories; G-code baselines are estimated from other participants. Lines and dots show participant means; diamonds and bars show group means and t-based 95 % confidence intervals (N = 14). The held-out low-AM versus white-noise difference is not reliable (Friedman p = 0.692; paired Wilcoxon p = 0.391). Panel A is exploratory and should not be read as a primary AM effect.

### Figure S3. High-Jerk Event Analysis

We found no evidence that high-jerk events increased lag-aligned tracking error relative to matched low-jerk pseudo-events (high-minus-low = 0.003 m; W = 67, p = 0.196; Figure S3). These results suggest that high source jerk does not trigger an all-or-none tracking failure.

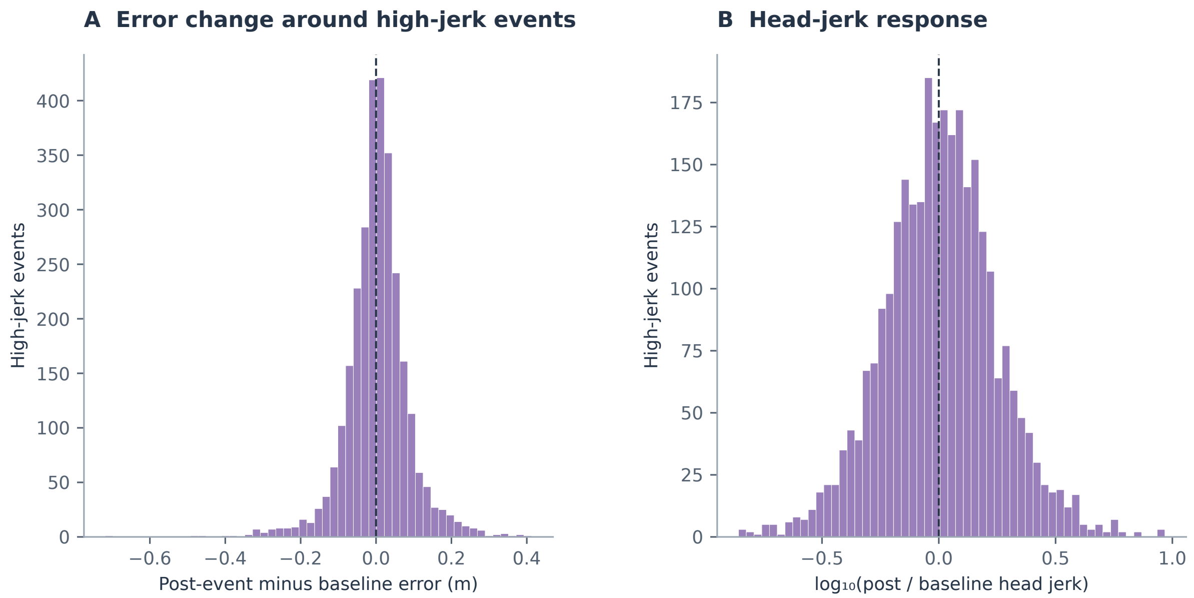


Figure S3: **Event-level responses around high source jerk.** **(A)** Distribution of post-event lag-aligned residual error minus local baseline. **(B)** $\text{Log}_{10}$ ratio of post-event to baseline head jerk; zero denotes equal jerk. Histograms contain 2,910 high-jerk events from auditory trials and are descriptive; events are not independent units of inference. The participant-level comparison of high- versus matched low-jerk pseudo-events showed no reliable increase in residual error (mean difference 0.003 m; $W = 67$, $p = 0.196$; $N = 14$). Post-event and baseline windows are defined in STAR Methods.

### Figure S4. DFC Delayed-Feedback Controller

As a complementary phenomenological model, head position was fitted with a point-mass PD controller with delay. Lateral median $R^2 = 0.56$; median $\zeta = 4.0$ (overdamped); median delay 0.60 s. No condition effects were found (all $F(3,39) < 2.5$, $p > 0.08$). The DFC provides a mechanistic interpretation of the tracking lag and overdamped kinetics, consistent with the delayed-state hybrid's long processing delay.

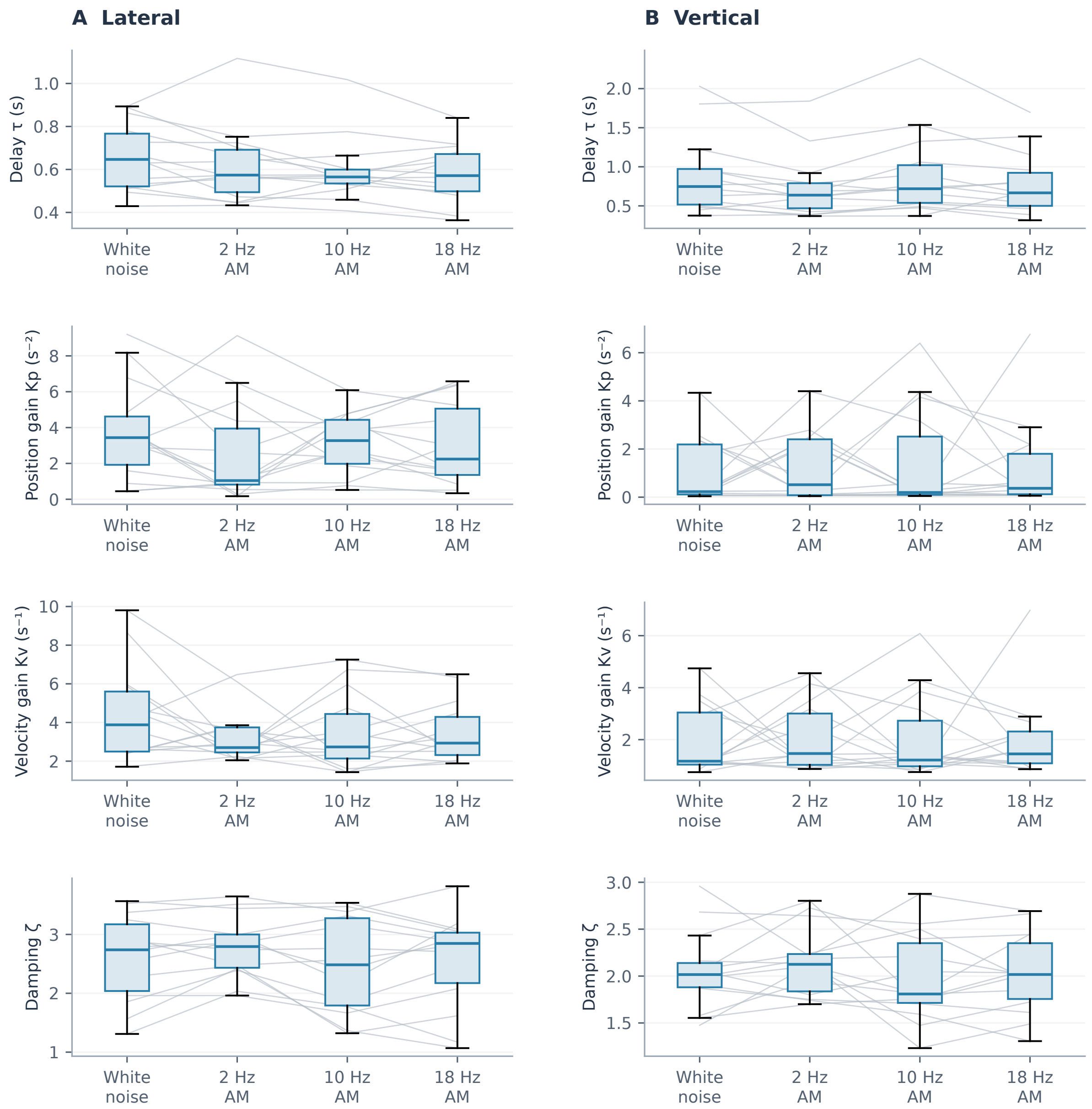


Figure S4: **Delayed-feedback controller parameters by auditory condition.** Lateral (A) and vertical (B) estimates are shown in separate columns; rows show delay $\tau$, position gain $K_p$, velocity gain $K_v$ and damping ratio $\zeta$. Each grey line connects one participant's mean across trials in the four conditions. Boxes show the median and interquartile range across participants; whiskers extend to 1.5 IQR, and outliers are omitted from the box display. The analysis includes 14 participants and excludes the visual familiarisation trials and participant A.P. Damping values near the fitting bound should be interpreted in light of the model constraints.

### Supplementary Note S1. Jerk-dependent gain extension

A jerk-dependent-gain extension of the delayed hybrid model, in which the gain matrices G and K scaled multiplicatively with instantaneous source jerk, was evaluated under the same G-code-grouped cross-validation scheme. This extension improved held-out prediction accuracy by an additional 0.000216 $m^2/s^2$ beyond the delayed hybrid model with constant gains (raw $p = 0.00043$; Holm-adjusted $p = 0.0017$ across four trajectory-feature gain tests). Extensions in which gains scaled with source speed, acceleration, or curvature did not survive Holm correction. The jerk-dependent-gain result is therefore the only trajectory-feature gain test that survives held-out cross-validation; the velocity-term share gradients of the delayed hybrid model in Figure S6 are descriptive covariation patterns rather than evidence for adaptive weighting. This represents a small improvement in a fitted gain-dependent extension and should not be interpreted as evidence of a neural gain-scheduling mechanism. The result is consistent with the absence of discrete jerk-triggered tracking failures (Figure S3): it reflects a modest predictive advantage of allowing gains to covary with source jerk, not a demonstrated behavioural adaptation to jerk events.

### Figure S5. Full model comparison

The per-participant held-out MSE for all five model variants under G-code-grouped five-fold cross-validation.

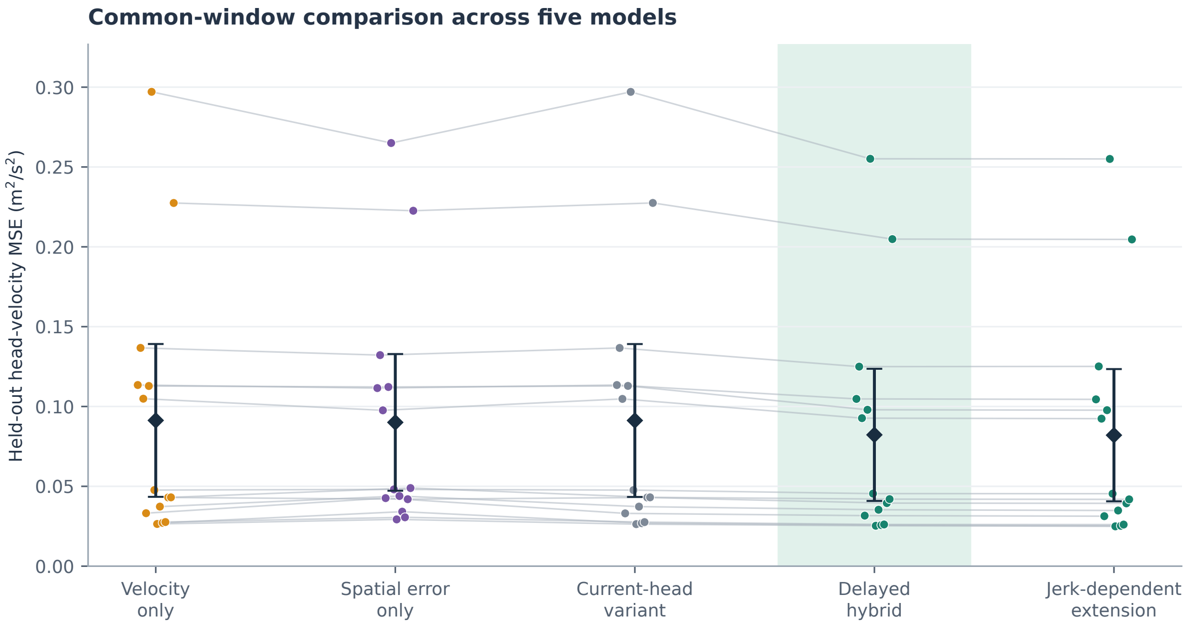


Figure S5: **Full model comparison on unseen trajectories.** Participant mean head-velocity MSE for the four primary models and the jerk-dependent gain extension, using G-code-grouped five-fold cross-validation and identical held-out samples for all models within a fold. Dots and connecting lines show 14 participants; diamonds and bars show the group mean and t-based 95 % confidence interval. The delayed hybrid model improves on each restricted model in all participants (one-sided $p = 6.1\times10^{-5}$). The jerk-dependent extension gives a further small mean MSE reduction of 0.000216 $m^2/s^2$ (raw one-sided $p = 0.00043$; Holm-adjusted $p = 0.0017$ across four feature-gain tests); this predictive improvement does not establish a neural gain-scheduling mechanism.

### Figure S6. Velocity-term share across all trajectory dynamics

Within-participant velocity-term share of the delayed hybrid model as a function of four trajectory features.

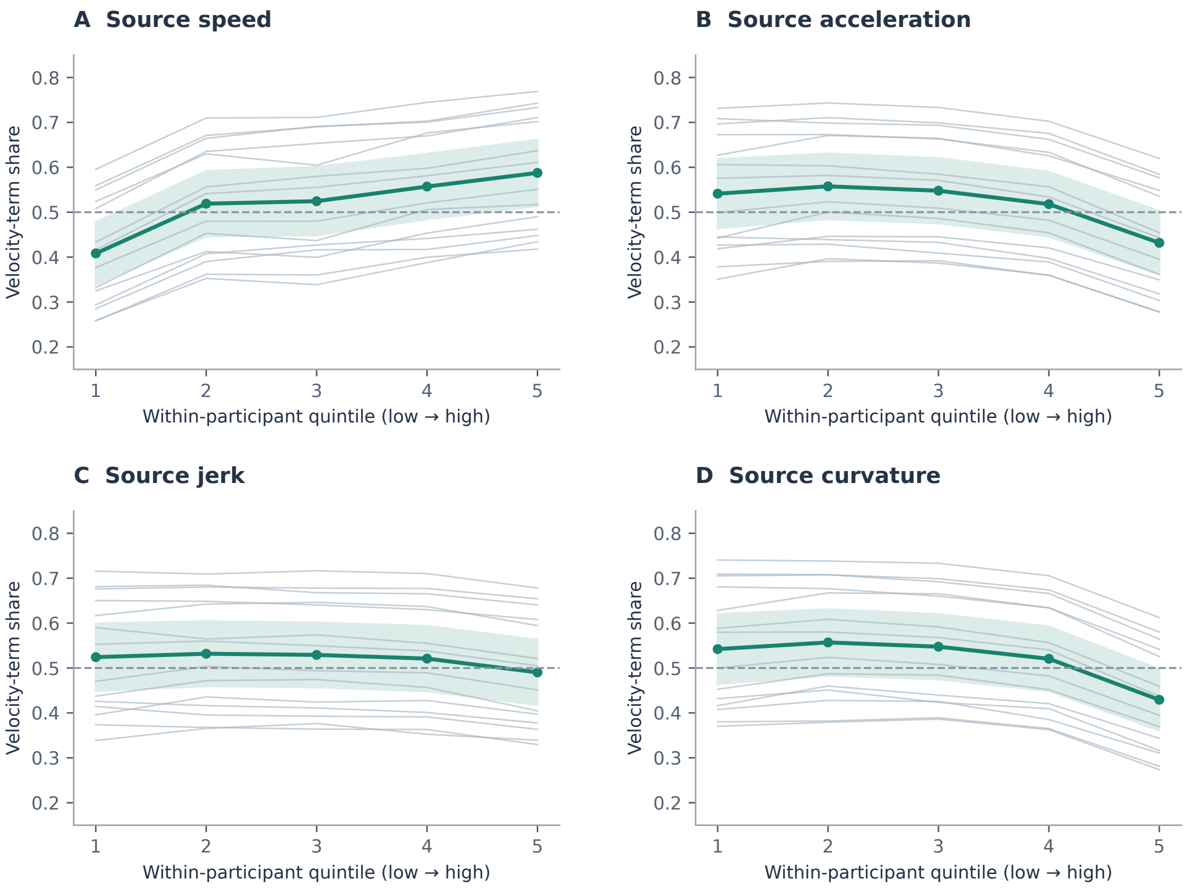


Figure S6: **Contributions of the two terms vary with trajectory statistics.** Panels show source speed (A), acceleration (B), jerk (C) and curvature (D). For each participant, samples are grouped into quintiles of the trajectory feature, and the velocity-term share is the magnitude of $G\dot{s}$ divided by the sum of the velocity- and spatial-error-term magnitudes, excluding the intercept. Thin lines show 14 participants; the green curve and ribbon show their mean and t-based 95 % confidence interval. The dashed line marks equal term magnitudes. This attribution analysis used trial-index-based folds rather than the G-code-grouped folds of the primary model comparison and is descriptive. The patterns reflect input covariation in the delayed hybrid model with constant gains, not adaptation of the gains to trajectory dynamics.
